# The Co-regulation Data Harvester for *Tetrahymena thermophila*: automated high-throughput gene annotation and functional inference in a microbial eukaryote

**DOI:** 10.1101/115816

**Authors:** Lev M. Tsypin, Aaron P. Turkewitz

## Abstract

Identifying co-regulated genes can provide a useful approach for defining pathway-specific machinery in an organism. To be efficient, this approach relies on thorough genome annotation, which is not available for most organisms with sequenced genomes. Studies in *Tetrahymena thermophila,* the most experimentally accessible ciliate, have generated a rich transcriptomic database covering many well-defined physiological states. Genes that are involved in the same pathway show significant co-regulation, and screens based on gene co-regulation have identified novel factors in specific pathways, for example in membrane trafficking. However, a limitation has been the relatively sparse annotation of the *Tetrahymena* genome, making it impractical to approach genome-wide analyses. We have therefore developed an efficient approach to analyze both co-regulation and gene annotation, called the Co-regulation Data Harvester (CDH). The CDH automates identification of co-regulated genes by accessing the *Tetrahymena* transcriptome database, determines their orthologs in other organisms via reciprocal BLAST searches, and collates the annotations of those orthologs' functions. Inferences drawn from the CDH reproduce and expand upon experimental findings in *Tetrahymena.* The CDH, which is freely available, represents a powerful new tool for analyzing cell biological pathways in *Tetrahymena.* Moreover, to the extent that genes and pathways are conserved between organisms, the inferences obtained via the CDH should be relevant, and can be explored, in many other systems.

## 1. Motivation and significance

*Tetrahymena thermophila* is a ciliate, one of the best-studied members of this large group of protists [1]. Its use as a model system led to the Nobel Prize-winning discoveries of telomerase and self-splicing RNA, as well as to other breakthroughs, including the isolation of dyneins and making the link between histone modification and transcriptional regulation [2, 3, 4, 5]. These contributions to our understanding of important cellular pathways made use of classical forward and reverse genetics, as well as biochemical approaches. More recently, genomic and transcriptomic data became available for *T. thermophila,* which have been used to infer functional gene networks [6, 7, 8, 9, 10, 11].

The *T. thermophila* genome has been sequenced and assembled [6], and is available online on the *Tetrahymena* Genome Database (TGD) [7]. While the TGD collates the sequence data along with available gene annotations and descriptions, the genome overall remains incompletely annotated. An extensive transcriptomic database, the *Tetrahymena* Functional Genomics Database or *Tetra*FGD, is also available for *T. thermophila* [10]. These data were collected over a well-established range of culture conditions in which *T. thermophila* undergoes large physiological changes [8, 9, 10, 11]. In addition to displaying individual expression profiles, the *Tetra* FGD can indicate the statistical strength of co-expression between any two genes, as calculated using the Context Likelihood of Relatedness (CLR) algorithm [9, 12, 13]. Co-expression in *T. thermophila,* as judged based on mRNA levels, can reveal functionally significant co-regulation. Gene regulation in this species appears to predominantly occur at the level of transcription [14], and so steady-state mRNA levels may explain the majority of steady-state protein levels, as reported in other systems [15]. In this report, we will refer to genes that are listed as co-expressed in the *Tetra* FGD as co-regulated.

A high-throughput analysis of *T. thermophila* gene expression profiles revealed that accurate gene networks can be inferred from co-regulation data [13], providing evidence that co-regulated genes tend to be functionally associated. This approach has been used in bacterial, mammalian, and apicomplexan systems [12, 16, 17]. There is also experimental evidence in *T. thermophila* to support the conclusion that co-regulation corresponds to functional association: Co-regulation data were used to successfully predict novel sorting factors and proteases involved in the biosynthesis of a class of secretory vesicles, called mucocysts [18, 19]. These results suggest that the *T. thermophila* transcriptome may be used to bioinformatically infer factors involved in an array of cellular pathways.

The CDH was designed to facilitate genome-wide analyses of gene co-regulation. The CDH automatically mines co-regulation data for genes of interest, and annotates the co-regulated genes *via* forward and reciprocal BLAST searches that identify orthologs in other model organisms. The CDH provides a systematic tool for gathering and annotating genomic information from public databases, and it can allow a researcher to quickly develop a robust hypothesis about the cellular pathways or structures in which a gene of interest may be acting, based upon the genes with which it is co-regulated.

## 2. Software description

### 2.1. Software Architecture

The CDH was developed for Python 2.7, along with the following packages: **sys**, **os**, **platform**, **logging**, **re**, **dill**, **difflib**, **csv**, **pdb**, **shutil**, **xml**, **win32com.shell**, **requests**, **BeautifulSoup4**, and **Biopython** [21].

Executables for Windows (x64) and MacOS (10.6+) were made using the Pyinstaller library. The CDH gathers available data for a set of co-regulated genes from publicly available databases, and uses these data to predict possible gene functions (Figure 1). The gathered information includes the co-regulation data from the TetraFGD, and the gene names, sequences, and annotations from the TGD. The available annotations come from a combination of experimental results and inferences from homology [7]. The CDH predicts annotations for genes based on the annotation of their respective orthologs, which are themselves identified by a series of forward and reciprocal BLAST searches *via* the National Center for Biotechnology Information (NCBI). These predicted annotations are generated by using the Ratcliff-Obershelp algorithm [22], as implemented in the python difflib library, to identify common phrases in the orthologs’ annotations.

**Figure 1:**
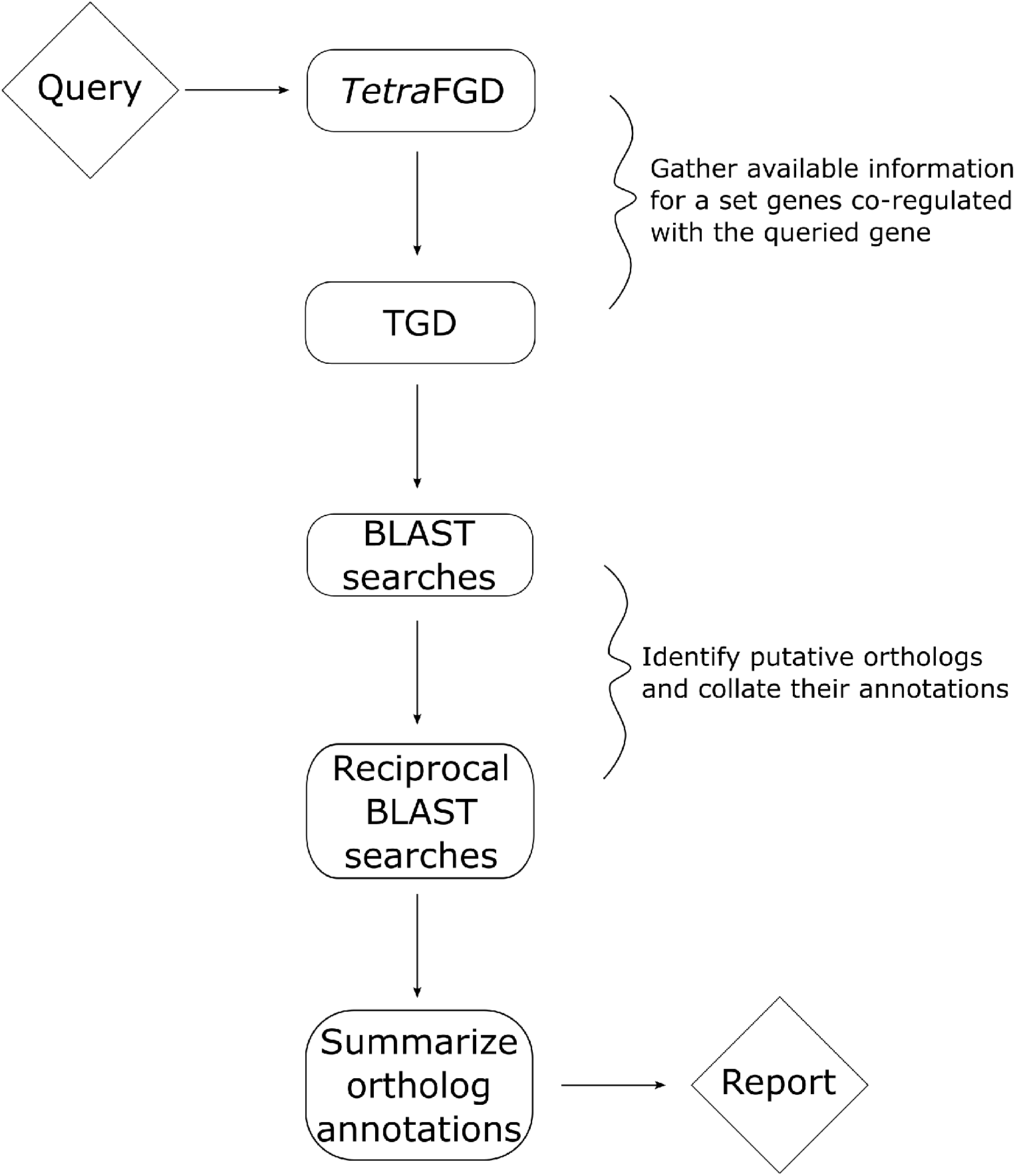
CDH architecture. Beginning with a single *T. thermophila* gene as a query, the CDH identifies all genes that are co-regulated with it, via the TetraFGD. Next, the CDH uses the TGD to gather the annotation and sequence data for each gene in the co-regulated set. For each gene in the co-regulated set, the CDH then runs forward and reciprocal BLAST searches, through the NCBI and TGD, to identify likely orthologs. A phrase matching algorithm, based on the Ratcliff-Obershelp algorithm [22], as implemented by the python difflib library, is then used to summarize the annotations of the putative orthologs for each *T. thermophila* gene in the co-regulated set. These summaries, which provide predictions about the function (e.g., relevant biological pathway) of the *T. thermophila* gene query, are presented along with the other data gathered, in the final report.

### 2.2. Software Functionalities

The basic functions of the CDH are to gather available co-regulation and annotation data, perform forward and reciprocal BLAST searches and predict gene annotations, and report this gathered information in a human-readable format. The CDH interface first asks the user to enter the ID for the gene whose co-regulated factors are of interest (Figure 2). Next, the user defines: how many of the co-regulated genes should be interrogated *via* BLAST; how to process data files that had been previously generated and are relevant to the current query; whether to use the BLASTp or BLASTx algorithm; and in which taxa to run the forward BLAST searches. The results of the CDH analysis are saved as a Comma Separated Values (.csv) file in the user's “Documents” folder: **/Documents/CoregulationDataHarvester/csvFiles**.

**Figure 2:**
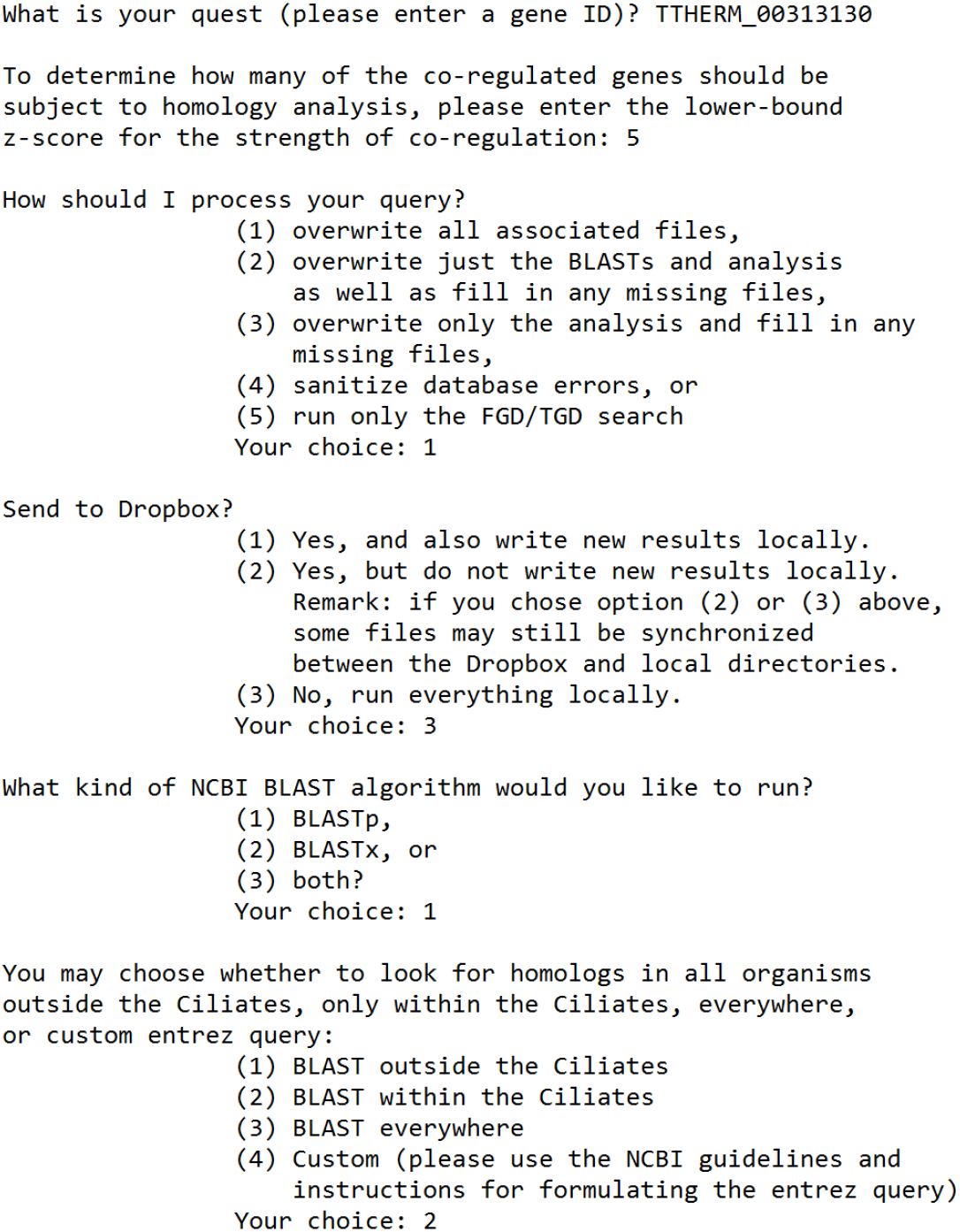
Setting CDH search parameters. The CDH is run through the terminal. The CDH prompts the user to define several parameters. These are: 1) the initial gene, i.e., the query; 2) the z-score threshold to be applied as cutoff for strength of co-regulation, which determines how many of the co-regulated genes will be subject to analysis *via* homology; 3) the extent to which data gathered in prior searches should be used; 4) whether results should be stored in Dropbox; 5) whether to run BLAST searches with cDNA or protein sequences; and 6) in which taxa to run the BLAST searches. For (2), the z-score threshold determines how many co-regulated genes will be included. For example, raising the threshold increases the stringency of the requirement for strength of co-regulation, so results in fewer co-regulated genes that are subsequently analyzed via BLAST, etc. For (3), the available options are: a) to run the search from scratch, overwriting any files associated with the queried gene; b) to re-use existing data for co-regulation, annotations, and sequences, but to run all of the BLAST searches from scratch; c) to re-use any existing data that are pertinent to the given query; d) to clear NCBI database errors from a previously run search and redo the associated BLAST searches; e) to only run the search for the co-regulation, annotation, and sequence. The example query in this screenshot is set to run a CDH search for the gene TTHERM_00313130 (Sortilin 4); to consider genes that are co-regulated with it with a z-score of 5 or higher; to gather all of the data de *novo;* to save all of the data locally; and to run the BLASTp searches only within the Ciliates.

## 3. Illustrative Examples

### 3.1. A CDH analysis of a factor required for programmed genome rearrangement returns the vast majority of experimentally-verified genes involved in the pathway

Programmed genome rearrangement is a tightly regulated process that occurs during the formation of the new somatic nucleus in conjugating *Tetrahymena* [23, 24]. This process is well-studied and known to be driven by a special adaptation of RNA interference, utilizing Dicer- and Piwi-like proteins, among other factors [25, 26]. *TWI1* encodes a Piwi-like protein that plays a central role in programmed genome rearrangement [27, 26]. When *TWI1* is entered as the query for the CDH, the CDH retrieves a large number of DNA and RNA-processing factors, as well as chromodomain proteins (Supplementary File 1). Importantly, these include the key factors known to be involved in programmed genome rearrangement (Table 1). The CDH report for this *TWI1* query is attached as Supplementary File 2. Within this report, we have highlighted the cases in which the CDH matched or expanded upon existing annotations.

It is also notable that specific homologs of Dicer that are not involved in programmed genome rearrangement, namely *DCR1* and *DCR2* [26], are not present in the list of genes co-regulated with *TWI1.* Similarly, while *TPB2* is a known genome rearrangement factor and is present in the CDH output [26], its paralog *TPB1* is neither involved in this process nor identified as co-regulated with *TWI1.* Thus, the CDH is a useful tool for focusing on pathway-specific paralogs within gene families.

**Table 1.** A subset of the programmed genome rearrangement factors that were identified by a CDH query for *TWI1*.

| Gene Group | Genes | Experimental Reports in *T. thermophila* |
| --- | --- | --- |
| Chromo-domain | *PDD1*,<br>*PDD2*,<br>*PDD3* | *PDD1* and *PDD2* are essential for programmed genome rearrangement; *PDD3* is reported to co-localize with *PDD1*, *PDD2*, and other necessary factors [32, 33, 34]. |
| Dicer-like | *DCL1* | *DCL1* is essential for programmed genome rearrangement [35, 36]. |
| Piwi-Associated | *GIW1* | *GIW1* is essential for programmed genome rearrangement and has no known conserved function [37]. |
| Localized in nuclear anlagen | *LIA1*,<br>*LIA2*,<br>*LIA3*,<br>*LIA4*,<br>*LIA5*,<br>*LIA6* | *LIA* proteins co-localize with *PDD1* at DNA rearrangement foci; *LIA1* and *LIA5* have been shown to be necessary for genome rearrangement, and *LIA3* is required for precise excision of eliminated sequences [38, 39, 40, 41]. |
| Zinc knuckle | *cnjB* | A double-knockout of *cnjB* and *WAG1* inhibits the formation of DNA elimination structures [42]. |
| Nucleic acid helicase | *EMA1* | *EMA1* is necessary for the association of *TWI1* with chromatin [43]. |
| Transposase | *TPB2* | *TPB2* catalyzes DNA elimination [44]. |
| Ku70/Ku80 beta-barrel domain | *TKU80* | *TKU80* is necessary for both *PDD1* complex assembly and DNA repair after programmed DNA elimination [45, 26] |
| Histone lysine methyl transferase | *EZL1* | *EZL1* is necessary for H3K27 methylation and programmed DNA elimination [46]. |

### 3.2. A CDH analysis of a mucocyst biogenesis factor enriches for mucocyst cargo and maturation factors

Mucocysts in *Tetrahymena* are secretory organelles. Mucocysts undergo a maturation process that requires the catalyzed cleavage of cargo proteins, called GRLs [28]. The *T. thermophila* genome encodes approximately 480 predicted proteases [6], but only five of these are co-regulated with GRLs, as revealed by a manual inspection of expression profiles on the *Tetra*FGD [19]. Two of these proteases, called *CTH3* (cathepsin 3) and *CTH4* (cathepsin 4), were subsequently shown to represent key enzymes for GRL cleavage [19, 29].

Using *CTH3* as a query for the CDH results in a list that includes a large number of genes known to be involved in mucocyst biogenesis (Table 2), and is enriched in membrane-trafficking factors and proteins with as-yet unknown functions in this organism (Supplementary File 3). Among the latter are a subunit of the *AP3* complex and a syntaxin in the *STX7* subfamily. Subsequent functional analysis of these genes showed that they are both essential for mucocyst formation, providing the best evidence to date that mucocysts are lysosome-related organelles (Kaur et al., submitted). The CDH report for this *CTH3* query is attached as Supplementary File 4. This report is also edited to indicate the cases when the CDH matched or expanded upon existing gene annotations.

**Table 2:** : Mucocyst biogenesis and cargo factors that were identified by a CDH query for *CTH3*.

| Gene Group | Genes | Experimental Reports in *T. thermophila* |
| --- | --- | --- |
| Sortilins | *SOR1*, *SOR2*, *SOR4* | *SOR2* and *SOR4* are essential for mucocyst biogenesis [18]. |
| Cathepsins | *CTH1*, *CTH2*, *CTH4* | *CTH1* and *CTH2* are involved in mucocyst biogenesis [19]. |
| Granule cargo | *GRL1*, *GRL2*, *GRL3*, *GRL4*, *GRL6*, *GRL7*, *GRL9*, *GRT1*, *IGR6*, *IGR7* | These genes, belonging to two gene families, encode mucocyst contents [28]. |
| Adaptin complex subunits | *AP3* $\mu$ and $\beta$ subunits | *AP3* $\mu$ subunit (*APM3*) is necessary for mucocyst biogenesis (Kaur et al., submitted). |
| Syntaxins | *STX7L1* | *STX7L1* is essential for mucocyst biogenesis (Kaur et al., submitted). |

## 4. Impact

The CDH reproduces existing annotations with high accuracy, and provides a large number of new annotations and expansions upon existing ones (Table 3; Supplementary Files 2 and 4). Effectively, the CDH increased the annotation coverage of the genes co-regulated with *TWI1* from 46% to 60%, and the annotation coverage of the genes co-regulated with *CTH3* from 41% to 57%. Specifying the BLAST parameters allows the user to discover the most informative functional predictions for their genes and pathways of interest. Limiting the CDH search to lineages outside of the ciliates is more likely to retrieve previously annotated orthologs, but runs the increased risk that weak homologs will generate spurious results. For some processes, such as programmed genome rearrangement in which *TWI1* is involved, the most informative BLAST searches may be those restricted to the ciliates. In our trials, the effectiveness of the CDH is maintained regardless of which taxa the BLAST searches are run against.

**Table 3:** The CDH accurately reproduces existing annotations and provides new annotations at a high rate.

| Queried Gene | *TWI1* | *CTH3* |
| --- | --- | --- |
| Number co-regulated genes | 932 | 213 |
| Taxa BLASTed | Only Ciliates | Excluding Ciliates |
| Number previously annotated | 430 | 88 |
| Annotations matched | 299 | 43 |
| Annotations expanded upon | 19 | 19 |
| Novel annotations | 126 | 34 |

**Table 4:**
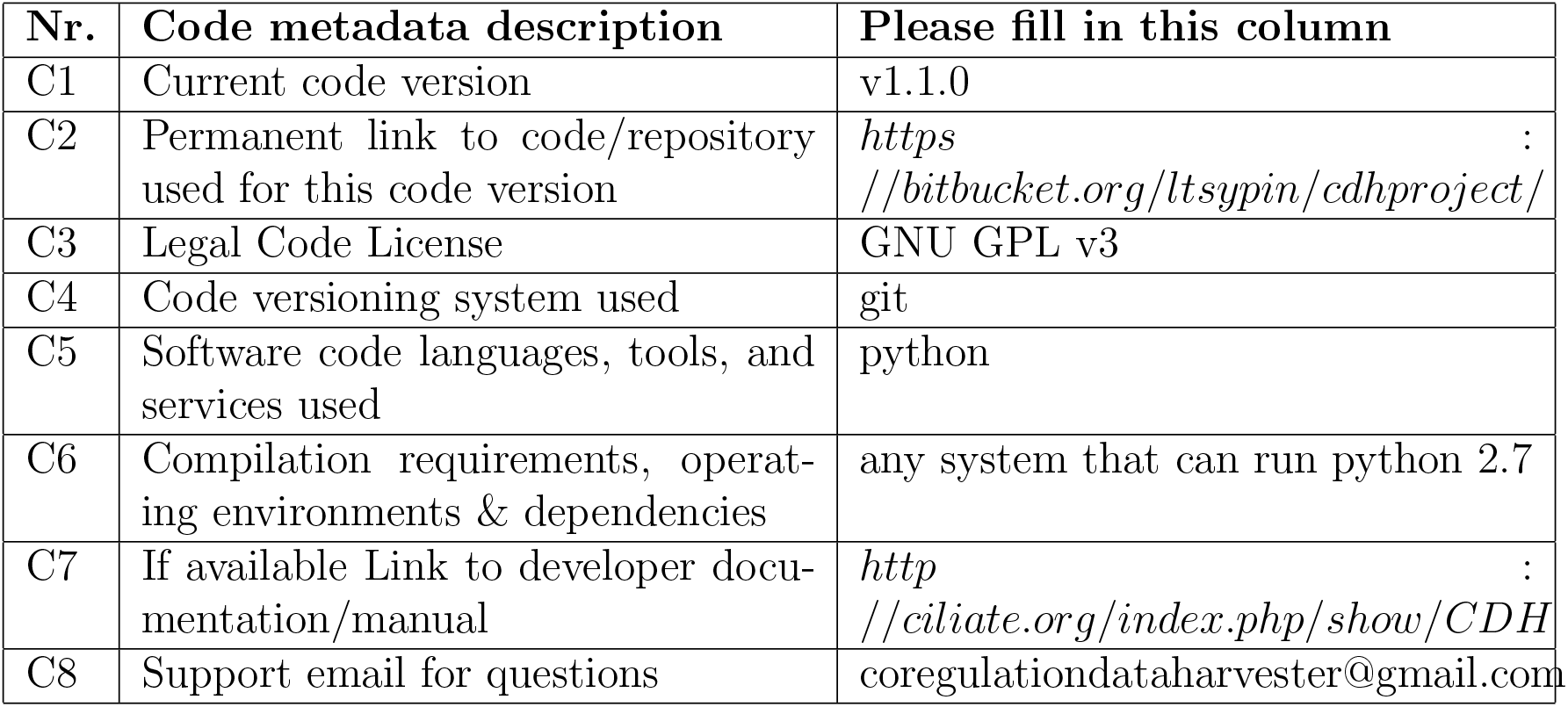
Code metadata (mandatory)

| Nr. | Code metadata description | Please fill in this column |
| --- | --- | --- |
| C1 | Current code version | v1.1.0 |
| C2 | Permanent link to code/repository used for this code version | https://bitbucket.org/ltsypin/cdhproject/ |
| C3 | Legal Code License | GNU GPL v3 |
| C4 | Code versioning system used | git |
| C5 | Software code languages, tools, and services used | python |
| C6 | Compilation requirements, operating environments & dependencies | any system that can run python 2.7 |
| C7 | If available Link to developer documentation/manual | http://ciliate.org/index.php/show/CDH |
| C8 | Support email for questions |  |

**Table 5:** Software metadata (optional)

| Nr. | (Executable) software metadata description | Please fill in this column |
| --- | --- | --- |
| S1 | Current software version | 1.1.0 |
| S2 | Permanent link to executables of this version | http://ciliate.org/index.php/show/CDH |
| S3 | Legal Software License | GNU GPL v3 |
| S4 | Computing platforms/Operating Systems | OS X (10.6+) and Windows (x64) Vista or later |
| S5 | Installation requirements & dependencies |  |
| S6 | If available, link to user manual - if formally published include a reference to the publication in the reference list | http://ciliate.org/index.php/show/CDH |
| S7 | Support email for questions |  |

In addition to providing a means of quickly gathering available data about a set of co-regulated genes and inferring their functions, the CDH data can be extended to to investigate the potential overlap between components of different cellular pathways. For example, *NUP50* encodes a gene that functions both in nuclear import at the nuclear pore complex and as part of a complex involved in transcription [30, 31]. Accordingly, the genes co-regulated with *NUP50* show extensive overlap with genes co-regulated with an import factor (Importin*β*) and with a gene involved in transcription (*RPB81,* an RNA polymerase II subunit), among other factors involved in both processes (Figure 3, A).

**Figure 3:**
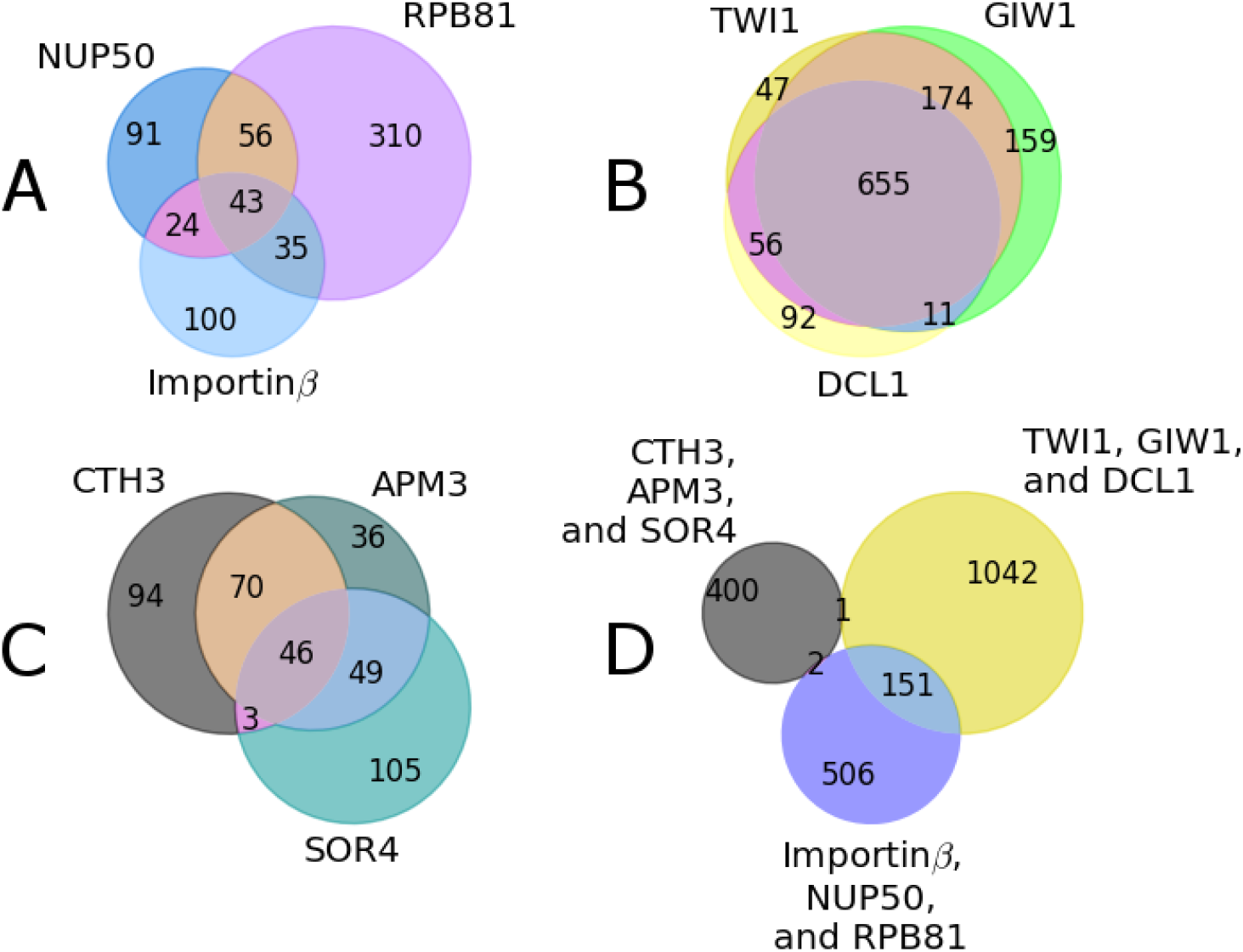
Using CDH outputs to assess overlap in gene function. Panels A, B, and C illustrate the overlaps in co-regulated genes for three different cellular pathways: A) nuclear import and transcriptional regulation; B) programmed genome rearrangement during cell conjugation; and C) mucocyst biogenesis. Each circle in the Venn diagrams corresponds to the full set of genes, as reported by the *Tetra*FGD, that are co-regulated with the gene indicated at the periphery of the circle. (A) *NUP50* (Nucleoporin 50) plays roles both in nuclear import and in gene transcription. The dual role of *NUP50* is reflected in the overlap of genes co-regulated with Importin β (an import factor) and with *RPB81* (RNA Pol II subunit), a transcription factor. *NUP50, RPB81,* and Importin β are mutually coregulated. The CDH identifies 214 genes co-regulated with the nucleoporin *NUP50,* 444 genes co-regulated with *RPB81,* and 200 genes co-regulated with Importin β. (B) *TWI1* (Tetrahymena Piwi 1), *GIW1* (Gentleman in Waiting 1) and *DCL1* (Dicer-like 1) are all required for programmed genome rearrangement, and are mutually co-regulated. The CDH identifies 932 genes co-regulated with *TWI1,* 999 genes co-regulated with *GIW1,* and 814 genes co-regulated with *DCL1.* (C) *CTH3* (cathepsin 3), *APM3* (μ subunit of the adaptin 3 complex), and *SOR4* (sortilin 4) are all required for formation of mucocysts, and are mutually co-regulated. These genes also appear to have distinct cellular functions in addition to their roles mucocyst formation. For example, mucocysts are non-essential organelles, yet *CTH3* is an essential gene. The CDH identifies 213 genes co-regulated with *CTH3,* 201 genes co-regulated with *APM3,* and 203 genes co-regulated with *SOR4.* (D) Pooling all of the genes represented in A, B, and C demonstrates that there is no overlap in co-regulated genes between A and B or C, and limited overlap between B and C.

It is informative to compare the overlap of co-regulated genes in different pathways. The co-regulated gene sets for three factors involved in genome rearrangement, *TWI1, GIW1,* and *DCL1,* show almost complete overlap, suggesting that they may be involved in a single common process (Figure 3, B). In contrast to the case of programmed genome rearrangement, the co-regulated gene sets for three factors required in mucocyst formation, *CTH3, SOR4,* and *APM3,* show partial overlap, hinting that one or more of these factors may also play roles unrelated to mucocysts (Figure 3, C). Consistent with this idea, *CTH3* is an essential gene, while mucocysts themselves are dispensable for cell viability in the laboratory [19]. Importantly, there is very little overlap between the co-regulated gene sets defined for the three different cellular processes (nuclear import/transcription, genome remodeling, and mucocyst formation) (Figure 3, D). The overlap is smallest between the genes co-regulated with mucocyst biogenesis factors and genes co-regulated with either nuclear import, transcriptional regulation, or programmed genome rearrangement. The somewhat greater sharing of genes co-regulated with nuclear import, transcriptional regulation, and programmed genome rearrangement may reflect the fact that these pathways all take place in the nucleus and are intrinsically linked to the cell cycle. Given the ease of assembling sets of co-regulated genes using the CDH, this type of overlap analysis can be extended to many cellular pathways.

## 5. Conclusions

Protists constitute the majority of eukaryotic diversity, meaning that this group needs to be included in evolutionary analyses of cellular processes, but this diversity is largely overlooked in the standard collection of model eukaryotes [20]. We present the Co-regulation Data Harvester for *T. thermophila* (CDH) as a tool that expedites analyses of *T. thermophila* genome, transcriptome, and cellular biology in an evolutionary context. The CDH is freely available and provides a systematic framework for genome annotation. It quickly gathers information from disparate databases and, by optionally reusing BLAST results that had been stored during previous queries, can increase in speed with successive uses. In providing a new means to analyze transcriptomic data, the CDH makes clear the potential for using the rapidly growing amount of genomic and transcriptomic data in many organisms, to facilitate functional analysis in poorly annotated or emerging model systems.

Users of the CDH should keep in mind that its reports are necessarily limited by pre-existing data from the TGD, *Tetra*FGD, and the NCBI. For example, the *Tetra* FGD does not provide co-expression data for genes whose expression level falls below a set threshold. Because of this limit, some *T. thermophila* genes may be overlooked by the CDH. Executable files for the program can be found at http://ciliate.org/index.php/show/CDH. A manual with detailed instructions and usage examples is provided in Supplementary File 5.

## Acknowledgements

We would like to thank Stefano Allesina for providing the initial impetus for this project, excellent advice, and review of this manuscript. Thanks to Chad Pearson and Alexander Stemm-Wolf for their feedback on the CDH and this manuscript. We thank Naomi Stover and Wei Miao for granting the Ciliate community access to the TGD and *Tetra*FGD, without which none of this work would be possible. We thank Tokuko Haraguchi for helpful discussion about NUP50. We thank Daniela Sparvoli and Harsimran Kaur for helpful discussion and good company. This work was supported by NIH GM-105783 to APT.

